# Evolutionary route of resistant genes in *Staphylococcus aureus*

**DOI:** 10.1101/762054

**Authors:** Jiffy John, Sinumol George, Sai Ravi Chandra Nori, Shijulal Nelson-Sathi

**Affiliations:** Computational Biology Laboratory, Inter Disciplinary Biology, Rajiv Gandhi Centre for Biotechnology (RGCB), Thiruvananthapuram, India; Manipal Academy of Higher Education (MAHE), Manipal, Karnataka, India

**Keywords:** Microbial genome evolution, Pan-genome, Antibiotic resistance, SCCmec, Lateral/Horizontal gene transfer

## Abstract

Multi-drug resistant *S. aureus* is a leading concern worldwide. Coagulase-Negative Staphylococci (CoNS) are claimed to be the reservoir and source of important resistant elements in *S. aureus*. However, the origin and evolutionary route of resistant genes in *S. aureus* are still remaining unknown. Here, we performed a detailed phylogenomic analysis of 152 completely sequenced *S. aureus* strains in comparison with 7,529 non-*S. aureus* reference bacterial genomes. Our results reveal<u>s</u> that *S. aureus* has a large open pan-genome where 97 (55%) of its known resistant related genes belonging to its accessory genome. Among these genes, 47 (27%) were located within the Staphylococcal Cassette Chromosome (SCCmec), a transposable element responsible for resistance against major classes of antibiotics including beta-lactams, macrolides and aminoglycosides. However, the physically linked mec-box genes (*MecA-MecR-MecI)* that are responsible for the maintenance of SCCmec elements is not unique to *S. aureus*, instead it is widely distributed within *Staphylococcaceae* family. The phyletic patterns of SCCmec encoded resistant genes in *Staphylococcus* species are significantly different from that of its core genes indicating frequent exchange of these genes between *Staphylococcus* species. Our in-depth analysis of SCCmec resistant gene phylogenies reveals that genes such as *blaZ*, *ble, kmA* and *tetK* that are responsible for beta-lactam, bleomycin, kanamycin and tetracycline resistance in *S. aureus* were laterally transferred from non-*Staphylococcus* sources. In addition, at least 11 non-SCCmec encoded resistant genes in *S. aureus*, mostly present in plasmid are laterally acquired from distantly related species. Our study evidently shows that gene transfers played a crucial role in shaping the evolution of antibiotic resistance in *S. aureus*.

## Introduction

Antibiotic resistant bacteria are causing serious global threat (World Health Organization 2016, Center for Disease Control 2013). Large population size and short generation time of bacteria are beneficial for the evolution of resistance within 2-4 years of introducing a new antibiotic (Harkins 2017). Clinically relevant opportunistic pathogens like *S. aureus* have acquired resistance against multiple antibiotics such as penicillin, methicillin, and vancomycin. Methicillin resistant *S. aureus* (MRSA) has become a major problem worldwide and is increasingly being detected in both hospitals and communities (Chambers and DeLeo 2009). The lateral component of microbial genome evolution (LGT/HGT) has played an essential role in shaping the bacterial genome content as well as their metabolic capabilities (Soucy SM, Huang J, and Gogarten JP 2015, Nelson-Sathi S. *et al.* 2015). For example, *S. aureus* metabolic capabilities are connected to the acquisition of both resistant and virulent traits (McCarthy *et al.* 2014, Bosi *et al.* 2016). About 15%–20% of the *S. aureus* genome includes parts of bacteriophage genomes, pathogenicity islands, plasmids, transposons, and cassette chromosomes (Lindsay 2010, Alibayov *et al.* 2014). The gene *MecA* is responsible for beta-lactam antibiotic resistance along with its regulatory proteins being embedded within the unique mobile genetic element called Staphylococcal Cassette Chromosome mec (SCCmec) (Ito *et al.* 2001). So far, 12 SCCmec types, designated as I to XII and different subtypes have been identified and the resistance capacity of the species varies based on its SCCmec types (Chongtrakool *et al.* 2006, Deurenberg and Stobberingh 2008). All SCCmec elements reported to date carry the following common gene complexes: (i) *orfX* (for the easy integration of SCCmec), (ii) five classes of mec-box genes (include *MecA/MecC/MecB* and its regulatory proteins *MecR* and *MecI* located upstream of *MecA*) (iii) flanking IS elements (*IS431* and *IS1272*) (iv) three allotypes of recombinases (*ccrA, ccrB*, and *ccrC;* sequence similarity between *ccr* allotypes is < 50%), which are responsible for the easy excision and integration of the SCCmec cassette (v) Transposons (*tnp* and *Tn554*, which are required for transposition) and (vi) other genes include heavy metal and antibiotic resistant genes such as *ars* and *blaZ* (IWG-SCC 2009, Ito *et al.* 2012).

Resistant genes encoded from different types and subtypes of highly variant SCCmec elements and other non-SCCmec regions; are the main reason behind the fast adaptation of *S. aureus* to different environments (IWG-SCC 2009). Previous studies on SCCmec elements have suggested that the *MecA* genes are distributed in multiple *Staphylococcus* species such as *S. epidermidis, S. haemolyticus* and *S. saprophyticus* (Grundmann *et al.* 2006). A recent study by Subakishita *et al.* found the presence of plasmid-encoded mec-gene complex (*mecAm-mecRm-mecIm-blaZm*) in a *Macrococcus caseolyticus* JSCS5402 strain, suggest that human pathogenic staphylococci acquired mec-box genes from *Macrococcus* lineage (Subakishita *et al*. 2010). Few other studies indicated that SCCmec elements assembled within Coagulase-Negative Staphylococci, such as *S. epidermidis, S. saprophyticus, S. schleiferi* and then transferred to *S. aureus* (International Working Group on the Classification of Staphylococcal Cassette Chromosome Elements IWG-SCC 2009), Ito *et al.* 2012, Otto 2013). In addition, study on *MecA* gene distribution pattern, Rolo *et al.* claimed that the most primitive staphylococcal species such as *S. fleurettii, S. sciuri*, and *S. vitulinus* were involved in the stepwise assembly of SCCmec elements and it is later transferred to *S. aureus* (Rolo *et al.* 2017).

However, there are limitations in the existing studies; mainly due to the lack of diverse data used. Most of these studies focused on SCCmec encoded elements, which contribute only 27% of resistance encoding genes in *S. aureus*. Many efforts have been taken to understand the evolution of antibiotic resistance in *S. aureus* and ended up with multiple observations, yet an overall picture is still lacking. Since most of the resistant genes in *S. aureus* located on mobile genetic elements and plasmids, we strongly hypothesize that gene exchange among closely as well as distantly related species, might have shaped the current form of resistant genes distribution in *S. aureus.* In the present study, we performed a detailed phylogenomic analysis of all the resistance encoding genes in 152 completely sequenced *S. aureus* strains and compared them with 7,529 reference bacterial genomes to construct a better picture of resistance evolution in *S. aureus*. Our findings reveal that the gene distribution and phyletic pattern of SCCmec elements in *Staphylococcus* species are significantly different from its core genes and which itself indicating that they are evolved in a different fashion. Our results show that LGT (Lateral Gene Transfers) plays a crucial role in shaping the evolution of SCCmec and non-SCCmec encoded resistant genes in *S. aureus*.

## Materials and Methods

### Data

Completely sequenced 7,681 bacterial genomes (protein sequences) and its feature tables were downloaded from National Center for Bioinformatics Information (NCBI) Genbank database (July 2017)(ftp://ftp.ncbi.nlm.nih.gov/genomes/genbank/bacteria/). This includes 152 *S. aureus* strains from different hosts (Human, Bovin, Chicken, Swine) isolated from different geographic regions and 7,529 bacterial reference genomes (Supplementary table 1).

### Pan-genome reconstruction and functional annotation of genes

Homologous genes were obtained from a total of 424,469 genes encoded in the chromosomes and plasmids of 152 *S. aureus* genomes. An all-against-all genomes BLASTp (Altschul *et al.* 1997) was performed with e-value <1e-05, ≥30% amino acid BLAST identity and ≥70% query coverage followed by reciprocal best BLAST hits (rBBH) (Tatusov *et al.* 1997). Homolog protein pairs were further globally aligned using Needleman-Wunsch algorithm with needle program of EMBOSS package (Rice *et al.* 2000) and homologs were filtered out with ≥25% global identity threshold. The resulting 428,070 protein pairs were clustered using the Markov Cluster algorithm (MCL) (Enright *et al.* 2002) with default parameters yielded a total of 4,565 protein families. There were 1,764 genes, which don’t have any significant homolog in 152 strains were classified as unique cases. 192 paralog genes, which skipped the MCL clustering pipeline, were manually added back to the existing clusters using a best cluster match criteria. Altogether 4,565 protein families and 1,764 unique genes were converted into a binary matrix of presence/absence patterns (PAPs). Based on the distribution pattern of clusters within *S. aureus* strains, its pan-genome is classified into three categories (genes present in >=90% strains as core genome, gene presence in >1%-<90% of strains as auxiliary genome and presence of a gene in a single strain as unique). The *S. aureus* pan-genome is summarized as a presence absence matrix where x-axis represents protein families and unique elements; where y-axis represents 152 *S. aureus* strains. If a protein family or unique gene is present in any of the *S. aureus* strain, its corresponding entry is represented with “1”; otherwise, a “0”. The presence absence matrix was sorted in ascending order of its distribution and visualized using MATLAB R2015a. Representative genes for each MCL cluster were functionally annotated using Clusters of Orthologus Groups (COG) database (Tatusov *et al.* 2003). Both protein families and unique genes were mapped to the COG database using BLAST and functional assignment was done. If a COG ID mapped to more than one category, the category R (general function prediction only) was assigned. The COG categories for each protein are given in supplementary table 2. Curve fitting of *S. aureus* pan-genome was performed using a power-law regression based on Heaps’ law n=*k*\*N^−α^, as described in Tettelin *et al.* 2005, 2008; Rasko *et al.* 2008. Fitting was conducted with the PanGP software (Zhao *et al*. 2014), where n is the expected number of genes for a given number of genomes, N is the number of genomes (i.e., 152), and *k* and α (α =1-γ) are free parameters that are determined empirically. According to Heap’s Law, when *α* > 1 (*γ* < 0), the pan-genome is considered to be closed and the addition of new genomes will not increase the number of new genes significantly. On the other hand, when *α* < 1 (0 < *γ* < 1), the pan-genome is open, and for each newly added genome, the number of genes will increase significantly (Tettelin *et al.* 2008).

### Identification of resistance related clusters (SCCmec encoded and non-SCCmec encoded)

Based on the classification proposed by International Working Group on Staphylococcal Cassette Chromosome (IWG-SCC 2009) 104 SCCmec encoded elements from different SCCmec typed reference strains and previously reported 173 plasmid/chromosome encoded resistant genes of *S. aureus* were downloaded from NCBI-Genbank database. Each reference element was blasted against 4,565 protein families and 1,764 unique genes and the best-hit entries (with an e-value <1e-05, query coverage ≥70 and BLAST identity ≥30) were considered as SCCmec elements and other resistant homologs. This yielded 47 SCCmec encoded and 130 non-SCCmec encoded multi drug resistant gene families in *S. aureus*. To identify the homologs of these resistant genes in non-*Staphylococcus aureus* species BLASTp was performed using previously described thresholds. Coverage matrix was constructed by calculating the proportion of genomes containing homologs within a non-*Staphylococcus* taxon. Distance between genomic locations of SCCmec homolog genes within a bacterial species was averaged to calculate the length of SCCmec elements.

### Classification of MRSA, MSSA and SCCmec typing

Reference *MecA* sequence was BLASTed against 152 *S. aureus* strains and based on presence or absence of homolog (e-value <1e-05, query coverage ≥70 and BLAST identity ≥90) strains were classified as resistant and sensitive. Similarly, *ccr* allotypes of known SCCmec reference strains were BLASTed and the presence or absence of homologs based on best hits (BLAST identity ≥80% and query coverage of ≥70%, e-value <1e-05), strains were classified into 12 SCCmec types (I to XII).

### Phylogenetic tree reconstruction and comparison

16S rRNAs, SCCmec and antibiotic resistant genes with their homologs were aligned separately using MAFFT (Katoh *et al.* 2002) with the options *–localpair, –maxiterate = 1000* and the alignment confidence score is evaluated using GUIDANCE2 server (Sela I., Ashkenazy H., Katoh K. and Pupko T. 2015). Maximum likelihood trees were reconstructed using RAxML (Stamatakis 2014) under the PROTGAMMAWAG model; with statistical support at nodes obtained by bootstrapping on 100 resample datasets. Maximum likelihood trees between Firmicutes and non-Firmicutes were re-rooted using the *nw_reroot* program of Newick Utilities (Junier and Zdobnov 2010) and visualized using FigTree version1.4.3 (Rambaut 2012). To compare the phyletic patterns of SCCmec, chromosome and core genes, we subsampled trees that contain at least 12 OTUs common in each set of trees (Supplementary table 3). Further, the pairwise distances between trees were computed by Robinson-Foulds (RF) metric (Robinson and Foulds 1981) using Phangorn package (Schliep KP 2011) and the Kernel Density Estimate (KDE) of the distances were plotted in R. Pairs of distance distributions were compared using a two-sample KS test. The distance comparisons were repeated by independently sampling a subset of trees in triplicates and RF distances were pooled together for the statistical analysis.

### LGT inference

Any topological discordance between gene trees and 16S rRNA reference tree at the species level is considered as potential LGTs. In 177 resistant gene trees, we condensed OTUs at species-level and compared the nearest neighbor relationship of each OTU with the reference tree (Supplementary table 4). A gene is considered as acquisition in *S. aureus* if it is absent in nearest neighbors and widely distributed in distantly related species, which are different from its reference phylogeny. Taxa that are branching within *S. aureus* clade are considered as exports from *S. aureus*. 16SrRNA phylogeny used for the comparison is provided in Figure 3 and its newick tree (including all OTUs) is provided in Supplementary table 4

## Results

### Characteristics of *S. aureus* pan-genome

A total of 428,070 genes encoded in the genomes of *S. aureus* strains were clustered into 4,565 protein families and 1,764 unique genes using Markov Chain Clustering procedure (Enright *et al.* 2002). The *S. aureus* pan-genome is open (α<1) and appears to be moderately expanding with the inclusion of new genomes (Figure 1, Figure S1). The openness of pan-genome and presence of mobile genetic elements in *S. aureus* is similar to many other bacterial species, for example *E. coli* that is known to have an essential capability of acquiring new genes into its genome via inter-species transfers (Touchon *et al.* 2009). Among 4,565 protein families and 1,764 unique genes in *S. aureus* pan-genome, 2,426 (38%) core genes are universally or nearly universally distributed (present in ≥90% strains) in 152 *S. aureus* strains (Figure 1, Supplementary table 2). Majority of these core genes are encoded for the basic machinery of cells including essential aspects of transcription and translation, which is similar to that observed in *Streptococcus* and *E. coli (*Lefébure and Stanhope 2007, Touchon *et al.* 2009). The auxiliary (2,136) and unique genome (1,764) of *S. aureus* was found to have a different functional distribution as compared to that of the core genome. Unlike unique and core genomes, the auxiliary genome of *S. aureus* is more involved in cellular processing and signaling functions, which is similar to other bacterial species (Ozer *et al.* 2014). Also, a large proportion (12%) of auxiliary and unique genes in *S. aureus* have been identified as mobile genetic elements like transposons and prophages genome (replication, recombination, and repair) (Supplementary table 2). *S. aureus* showed an extensive genetic diversity, with an average 27% of genes being specific for a single strain.

**Fig. 1.**
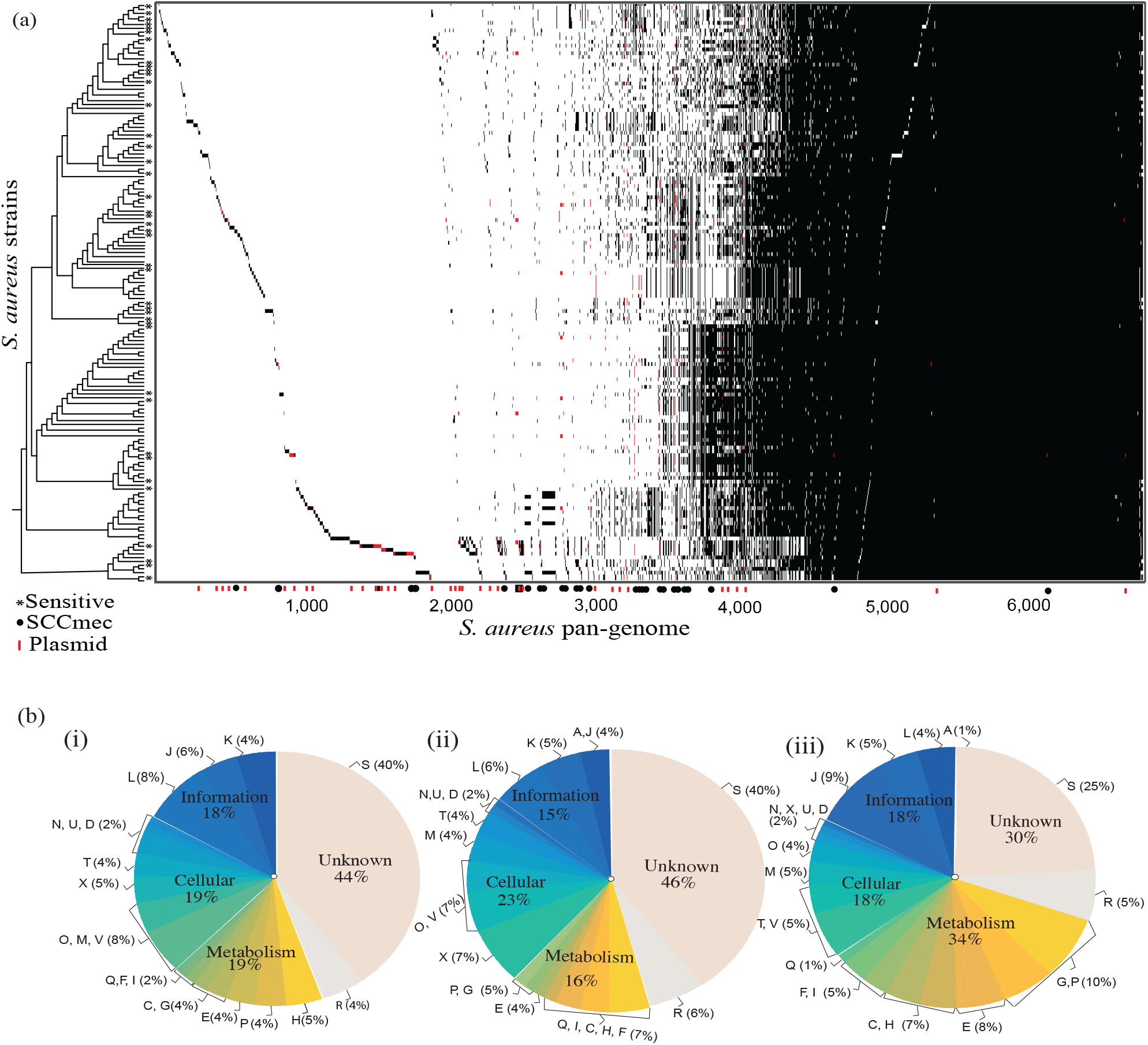
Pan-genome of *S. aureus*. (a) For each protein, a black tick indicates the presence and white tick indicates the absence of a gene in the corresponding genome in the left. *S. aureus* species on the left side are sorted according to the 16S rRNA reference tree. Genes encoded from plasmid are indicated in red in the presence and absence pattern. Methicillin susceptible *S. aureus (*MSSA) are marked with a * symbol. (b) Functional classification of *S*. *aureus* proteins using COG (i) the unique genome (genes that are unique to a single strain) (ii) the auxiliary genome (genes present in >1%<=90% of strains), and (iii) the core genome (genes present in ≥ 90% of strains). COG functional categories are as follows: For cellular processes and signaling: cell cycle control, cell division and chromosome partitioning (D); cell wall/membrane/envelope biogenesis (M); cell motility (N); mobilome, prophages, transposons (X); post translational modification, protein turnover, and chaperones (O); signal transduction mechanisms (T); intracellular trafficking, secretion, and vesicular transport (U); defense mechanisms (V); extracellular structures (W); nuclear structure (Y) and cytoskeleton (Z). For information storage and processing: RNA processing and modification (A); translation, ribosomal structure, and biogenesis (J); transcription (K) and replication, recombination, and repair (L). For metabolism: energy production and conversion (C); amino acid transport and metabolism (E); nucleotide transport and metabolism (F); carbohydrate transport and metabolism (G); coenzyme transport and metabolism (H); lipid transport and metabolism (I); inorganic ion transport and metabolism (P) and secondary metabolites biosynthesis, transport, and catabolism (Q). General function prediction (R).

### Evolution of SCCmec elements in *S. aureus*

Among 4,565 protein families and 1,764 unique genes, 47 protein families pertain to SCCmec encoded genes including *MecA* gene and its regulators (*MecR* and *MecI*), recombinases (ccrABC), insertion elements, transposons, heavy metal and drug resistant genes. Seventy-three percent of 152 *S. aureus* strains contains the *MecA* gene and were classified as ‘resistant’, while the remaining 27% strains were classified as susceptible. As expected, *MecA* was absent in most of the sensitive strains, but surprisingly 10% of sensitive *S. aureus* strains in our dataset consist of SCCmec elements without *MecA* gene, this suggests that the presence of SCCmec is not limited to resistant strains. This distribution can also be due to the loss of *MecA* gene after the acquisition of the well-established SCCmec elements. Furthermore, based on the *ccr* gene allotypes classification, all the 152 strains were found to be in well-established I to XII SCCmec types; with community-associated *S. aureus* (Type IV) being the most abundant among them (63%) (Supplementary table 1). As suggested by Berglund *et al.* in 2005, the observed wide distribution of Type IV *S. aureus* strains might be due to its small SCCmec size and the presence of the high number of functional recombinases. Among the 47 SCCmec encoded genes, the crucial genes responsible for the maintenance and functioning of the SCCmec elements (*MecA*, *MecR, MecI*, *orfX*, *IS431*, *Tn554A/B, tnp*, and some *ccr* allotypes) were nearly universally distributed among the selected 152 *S. aureus* strains (Figure 2, Figure S2). The cassette flanking gene viz. *orfX* was found to be universally (99%) present in *S. aureus* strains, clearly indicating its importance in the integration of these elements. The mec-box genes (*MecA-MecR-MecI*), which are the decisive elements in SCCmec, are predominantly distributed among 152 *S. aureus* strains (73%, 73%, and 64% distribution respectively). The ccr-box genes that code for recombinases are widely distributed within resistant strains of *S. aureus* (66%). Some of its allotypes such as *ccrAB4* and *ccrB1* occur at a lower frequency (1.3%) when compared to other *ccr* allotypes namely *ccrA1* and *ccrB2*. In addition to the mec-box and ccr-box genes, various IS elements and genes associated with the mobility of SCCmec were also found to be widely distributed within *S. aureus* (Figure 2). Moreover, heavy metal resistant genes such as *ars* (arsenate) and *cop* (copper) are also universally distributed in 152 *S. aureus* strains and their co-existence with the SCCmec elements hints the co-selection under environmental pressure (Xue *et al.* 2015). Interestingly, most of the SCCmec core elements (*orfX, MecA* and its regulators, *ccr* genes, IS elements, transposons, and heavy metal resistant genes such as *ars* and *cop*) are widely distributed within *Staphylococcaceae* with high sequence similarity (≥70% sequence similarity, Figure 2). While considering the entire Firmicutes phylum, the SCCmec homologs of *S. aureus* observed in many *Bacillales*, *Clostridiales*, and *Lactobacillales* species (Figure S2).

**Fig. 2.**
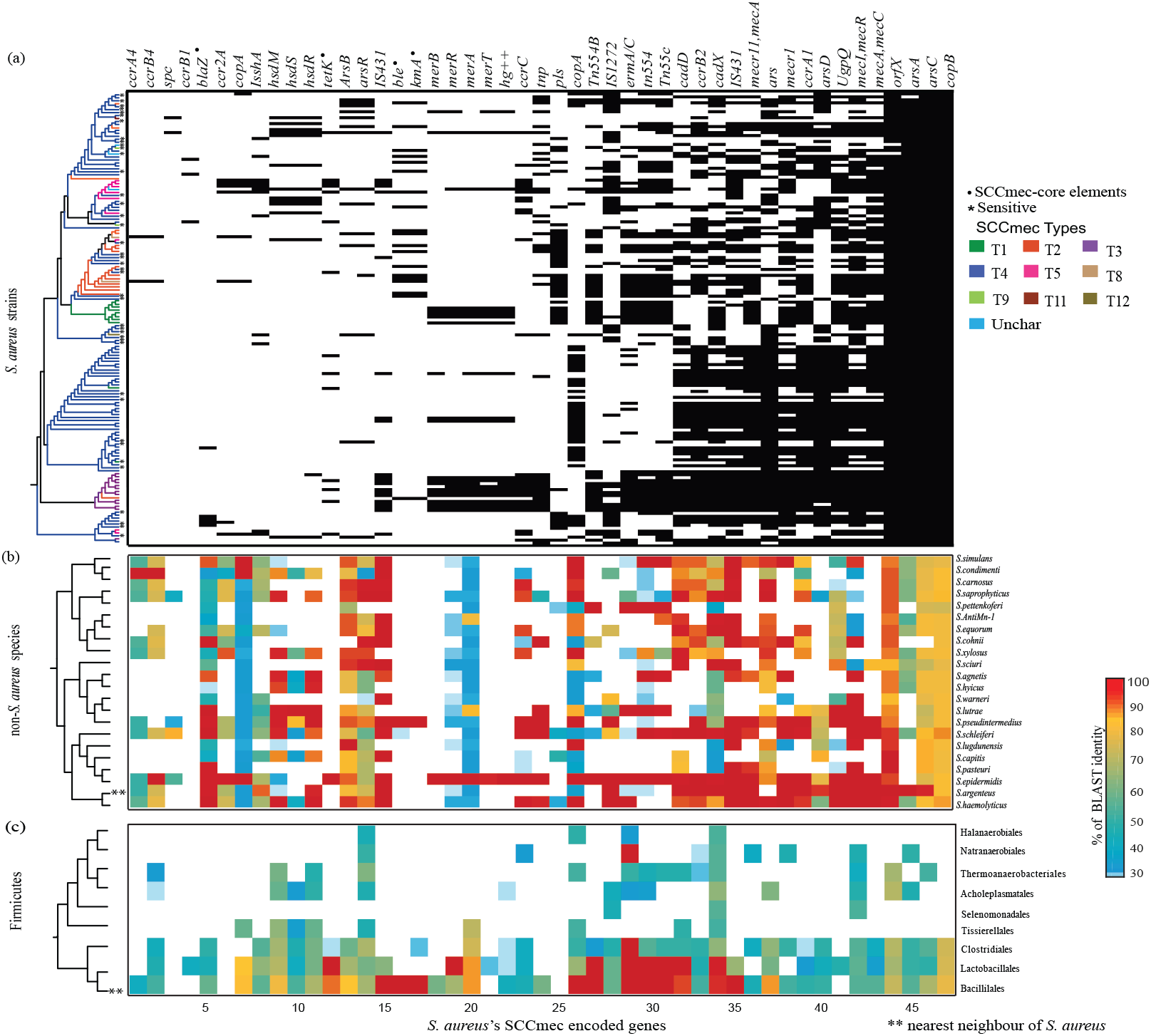
Distribution of 47 SCCmec encoded protein families in Firmicutes phylum. X-axis represents 47 SCCmec protein families and Y-axis represents 152 *S. aureus* strains, 22 non-*S. aureus* species and 9 orders within Firmicutes phylum. (a) The matrix indicates presence (black) or absence (white) of the SCCmec elements in the corresponding genome on the left. The 16S rRNA tree of 152 *S. aureus* strains is colored based on known 12 SCCmec types. Methicillin sensitive strains are indicated by an asterisk (*) symbol in the phylogenetic tree. LGT influenced genes are represented as a black dot (.). The *ccrA4* and *ccrB4* are patchily distributed because of the uniqueness in their sequence. (b) The matrix depicts the presence and similarity of 47 *S. aureus* SCCmec encoded genes in 22 non-*S. aureus* species and the color bar on the right represents sequence similarity ranging from 30 (blue) to 100 (red). (c) The matrix represents the presence of *S. aureus* SCCmec gene homologs in 9 orders within the Firmicutes phylum.

The physically linked core mec-box genes (*MecA-MecR-MecI*) ubiquitously present within *S. aureus* is necessary for the proper functioning of SCCmec cassette. The genomic organization of SCCmec-core genes, *MecA-MecR-MecI*, is highly conserved within resistant strains of *S. aureus* as well as in some of the *Staphylococcaceae* species including *S. sciuri, S. pseudointermedius, S. epidermidis, S. argenteus, S. haemolyticus, S. schleiferi* and *Macrococcus caseolyticus.* In these species, mec-box genes are located within 51 Kbps regions (Figure 3, Supplementary table 5), which is similar to those of SCCmec carried by *S. aureus* (Ito *et al.* 2012). In terms of sequence similarity and linkage, all the species mentioned above seems to have genes that are comparable to the *S. aureus* mec-box genes. On the other hand, some *Staphylococcus* species including *S. simulans, S. cohnii, S. lugdunensis, S. warneri, S. agnetis, S. saprophyticus*, and *S. equorum* have mec-box genes, but they are not located within the 51 Kbps range and don’t seem to have any apparent genomic linkage.

**Fig. 3.**
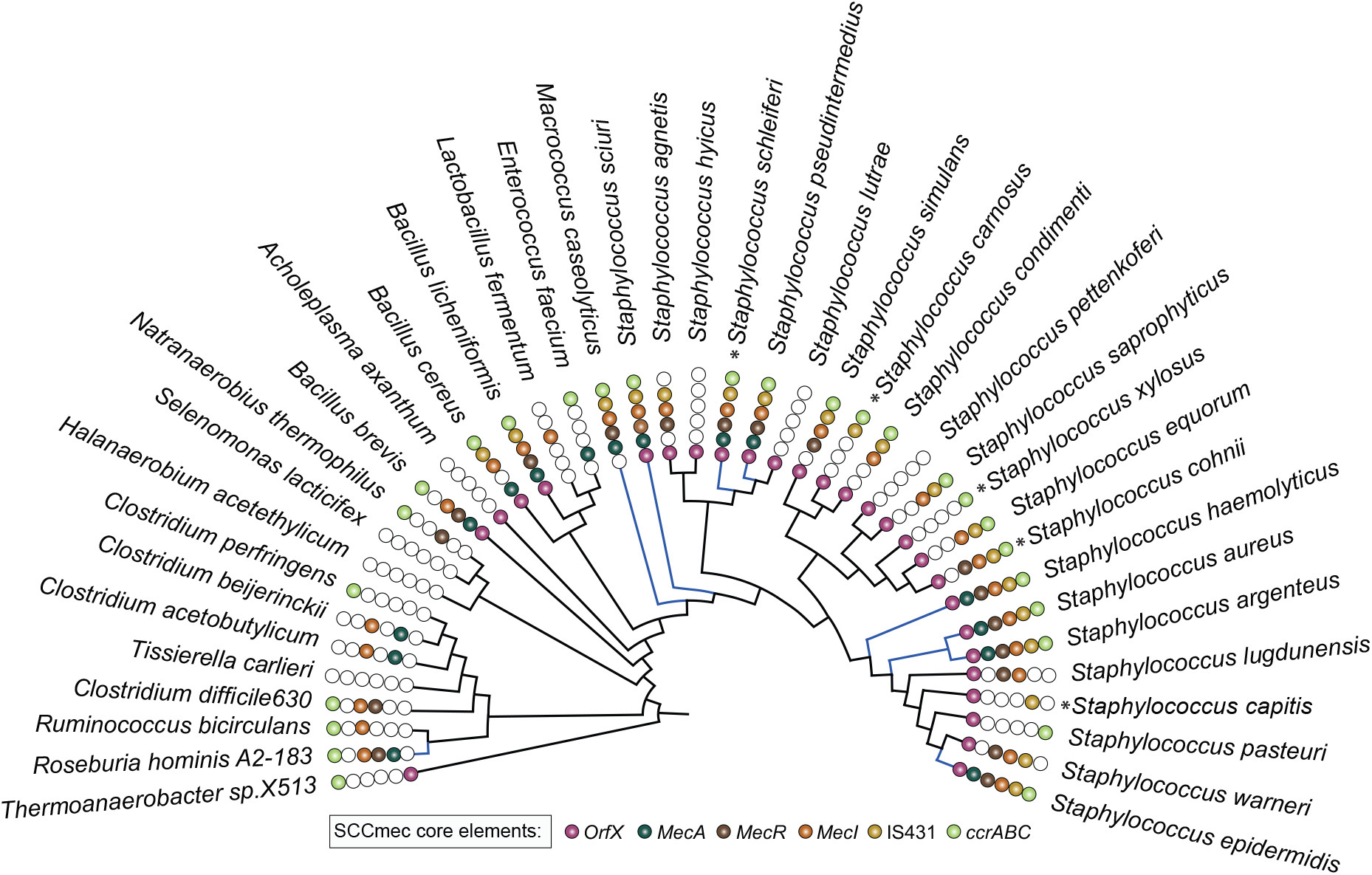
Distribution of SCCmec encoded genes within Firmicutes phylum represented in maximum likelihood phylogeny. Colored circles represent the presence of *orfX-mec* box*-ccr* box genes identified in different bacterial genera. Blue colored branches in the phylogeny represent the species with a genomic organization of the core SCCmec elements (*MecR-MecA-MecI*) within 51 Kbp genomic regions. Most of the species in *Staphylococcaceae* family are pathogens, while the opportunistic pathogens are represented as asterisk (*).

From gene distribution pattern and genomic linkage analysis it is clear that, SCCmec elements are widely present within *Staphylococcus* species. Moreover, the phylogenetic analysis of 47 genes encoded within the SCCmec cassette reveals that at least 42 (89%) of them are exhibiting significant topological discordances with respect to 16S rRNA reference phylogeny, while none of them are recovering individual species monophyly (Supplementary table 4). Tree dissimilarity distribution (Robinson-Foulds distances) of SCCmec and core genes present in *Staphylococcus* species is significantly different from each other (p-value < 2.2e-16) (Figure 4) indicating frequent exchange of these genes within the *Staphylococcus* species. Lateral transfers of SCCmec genes were not limited within *Staphylococcus* species; rather distantly related lineages shared genes with *Staphylococcus* species (Figure S3, Supplementary table 4). Resistant genes such as *blaZ* (beta-lactam resistance), *kmA* (kanamycin resistance), *tetK* (tetracycline resistance) and *ble* (bleomycin resistance) that embedded within the SCCmec elements of *S. aureus* are some of the good examples of laterally transferred genes from distantly related non-*Staphylococcus* species (Figure S4). For instance, *S. aureus* might receive *blaZ* gene from *Geobacillus stearothermophilus;* a species where there is no evidence of SCCmec cassette but phylogenetically they are showing a close relationship with *S. aureus*. Another example is *ble* in *S._aureus* that might have been acquired from *Paenibacillus beijingensis* species. In case of *tetK*, *S. aureus* received it from one of its distantly related lineages such as *Lactobacillales* and *Bacillales*. Furthermore, kanamycin resistant gene might have been acquired from *Paenibacillus riograndensis* species (Figure S4). These precisely indicate that lateral influence of closely and distantly related species largely contributed to the current structure of SCCmec cassette of *S. aureus*.

**Fig. 4.**
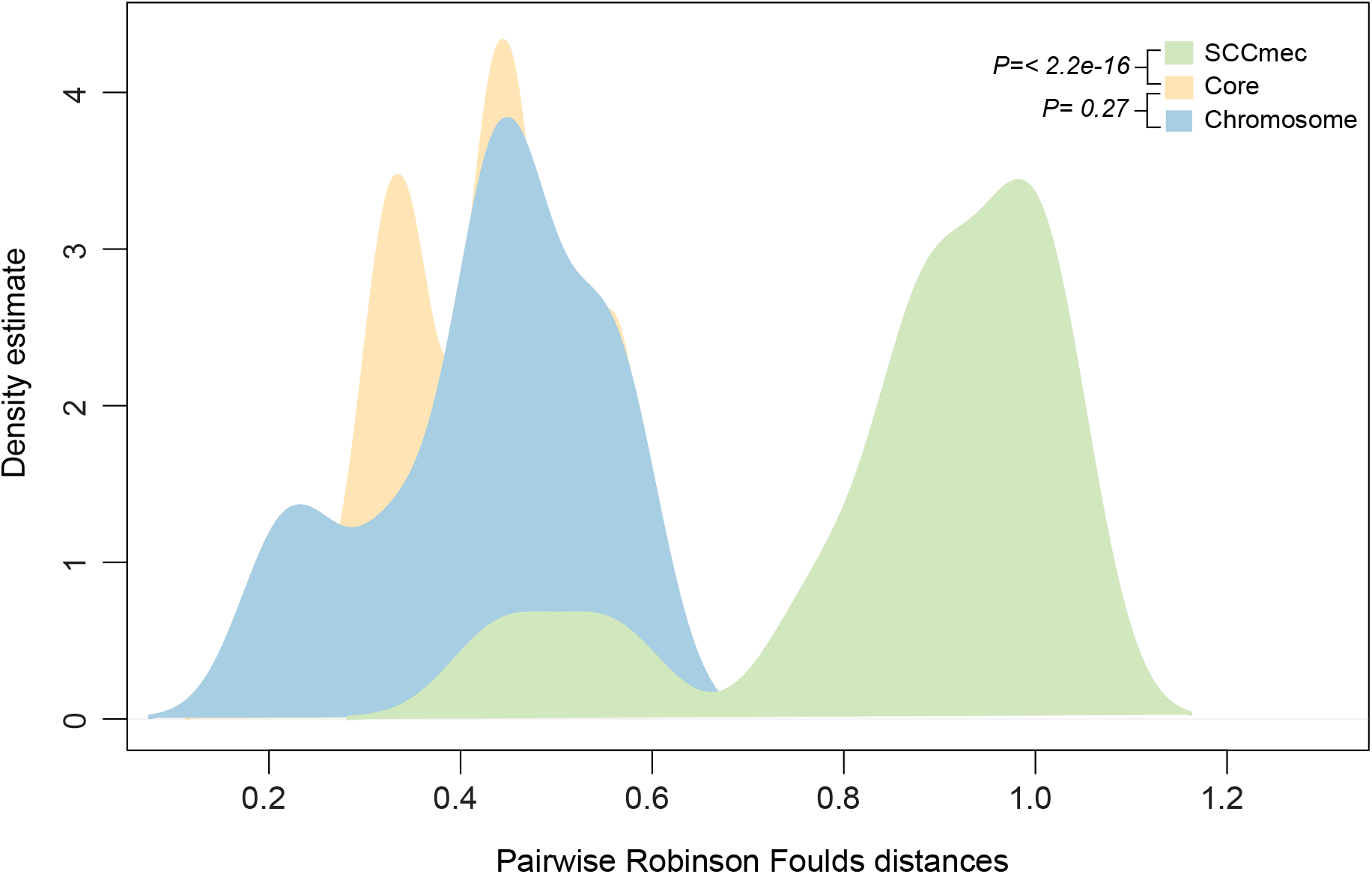
Distribution of pairwise tree distances of core and resistant genes. The pairwise tree distances of core, chromosome and SCCmec encoded resistant genes of *Staphylococcus* were calculated using Robinson-Foulds distance are shown. The X-axis represents the distances between gene trees and reference tree and the Y-axis represent their densities. P-values (two-tailed Kolmogorov-Smirnov test) from comparisons between core and SCCmec, core and chromosome genes were < 2.2e-16 and 0.27 respectively.

The *MecA* gene was subject to multiple lateral transfers within the Staphylococcace family members and Macrococcus *caseolyticus* species forming nearest neighbor to *S. aureus* (Supplementary table 4). In Macrococcus *caseolyticus* species, MecA gene is present in plasmid which increases the possibility that the MecA in *S. aureus* is originated from Macrococcus lineage. Moreover, species such as *S. sciuri, S. pseudointermedius, S. epidermidis, S. argenteus, S. haemolyticus* and *S. schleiferi* are branching within *S. aureus* clade pointing exports of *MecA* from *S. aureus*.

### Evolution of non-SCCmec encoded resistant genes

Even though SCCmec elements are known to be the key factors responsible for antibiotic resistance in *S. aureus*, there are about 130 non-SCCmec encoded resistance related genes encoded from plasmids and the chromosome of *S. aureus*. The majority of these genes confer resistance to protein synthesis inhibitor class of antibiotics (33%) while a small number are showing resistance to folic acid synthesis inhibitors (0.38%). Similar to the SCCmec encoded gene distribution, the non-SCCmec encoded resistant genes are also widely distributed within most of the *Staphylococcus* species (≥70% sequence similarity, Figure S5).

Among 130 non-SCCmec encoded resistant genes in *S. aureus* 83 are encoded from chromosome while 47 are from plasmid and more half of them showing topological resemblance with reference phylogeny. There are 53 genes showing *S. aureus* monophyly and nearest neighbor relationship as in 16S rRNA reference phylogeny (Supplementary table 4). Among these 53 genes, majority (87%) of them are chromosome-encoded, which suggest that a large fraction of vertically evolved genes retained within these species (Figure S6). Robinson-Foulds distance metric computed for all pairwise comparisons revealed that the phylogenetic distributions of chromosome encoded resistant genes are more similar to core genes (with p-value < 0.27) (Figure 4). Examples of vertically evolved resistant genes include *mgrA, arlS, murA, sulA* and *norB* that are responsible for resistance against different classes of antibiotics such as fluoroquinolone, cephalosporin, sulfonamide and fosfomycin. Even though a large proportion of chromosome encoded resistant genes seems to be evolved vertically within *Staphylococcus* species, lateral gene transfers also influenced genes from this category. There are few genes in *S. aureus* including *tetO, blmS, oppB, lnuA/linA and OppC* are examples of gene transfers from distinctly related linages (Figure S7a).

As expected, frequent LGTs were detected on plasmid-encoded genes in *S. aureus* which includes important genes such as *mupA, aadA5, tetL* that are responsible for resistance against mupirocin, aminoglycoside and tetracycline respectively (Figure S7b). Transfers from distantly related species such as *Clostridiales*, Proteobacteria and Actinobacteria lineages influenced most of the gene acquisitions. For example, aminoglycoside resistant gene *aadA5* phylogeny indicates that *S. aureus* might receive this gene from one of the distantly related lineages (*Clostridiales, Bacteroidetes* and *Virgibacillus*); but in case of *tetL*, *S. aureus* might have acquired it from one of the species of *Lactobacillales and Bacillales* (Figure S7b). Moreover, some genes such as *dfrA* and *mupA* were acquired it from the distantly related lineages in *Bacillales*.

## Conclusions

To better understand the evolution of antibiotic resistance-related genes in *S. aureus*, we have analyzed all known resistant genes present in 152 completely sequenced *S. aureus* genomes in comparison with 7,353 non-*S. aureus* reference bacterial genomes. The gene distribution pattern clearly shows that the SCCmec elements carry the antibiotic-resistant genes and its regulators are not unique to *S. aureus*. Moreover, they are widely distributed among other *Staphylococcus* species also present in some distantly related non-*Staphylococcus aureus* species. Furthermore, many *Staphylococcus* species were observed to have a physically linked mec-box-like structure in their genomes with a similar structural organization and high amino acid sequence identity to the SCCmec element of *S. aureus.* Bacterial species having such SCCmec element is patchily distributed within *Staphylococcus* species. Tree dissimilarity distributions of SCCmec genes in *Staphylococcus* species with its core genes indicate frequent exchange of genes encoded from this element happened within the *Staphylococcus* species. The lateral gene transfers were not limited within *Staphylococcus* species; rather unique resistant genes such as *blaZ*, *kmA*, and *ble are* acquired from distantly related lineages. In 130 non-SCCmec encoded resistant genes, most of the chromosome encoded resistant genes seem to be vertically inherited, while plasmid genes are more prone to lateral gene transfers. Our phylogenomic analysis provides a better insight into the evolution of resistant genes in *S. aureus* strains and role of LGTs shaping its current distribution.

## Discussion

For better insights on the resistance evolution in *S. aureus*, we have considered completely sequenced bacterial genomes (all publically available) in our analysis that is a limiting factor in the previous studies. The gene sharing patterns of *S. aureus* strains were not uniform between different resistant types; for example, Type IV strains are sharing more genes each other, but having less number of genes in its SCCmec element than others (Figure S1). Interestingly, we noticed that the 16S rRNA reference phylogeny of *S. aureus* strains shows no monophyletic structure in terms of its resistant gene presence or SCCmec type, indicating that *S. aureus* lineages were not primarily evolved toward the resistant trait (Figure 2a) and LGTs might play a crucial role in shaping its resistant genes distribution. Some previous reports show that SCC*mec* element is originated in Coagulase-Negative Staphylococci (CoNS) such as *S. epidermidis, S. haemolyticus, S. schleiferi*, and *S. sciuri* (Otto *et al.* 2013; Ito *et al.* 2012). However, we found that SCCmec elements are not limited to CoNS; rather mec-box and ccr-box gene homologs were observed in Coagulase-Positive Staphylococci such as *S. argenteus* and *S. pseudointermedius*. The physically linked mec-box genes (*MecA-MecR-MecI*) are widely distributed in *Staphylococcus* species, except in *Macrococcus caseolyticus* and *Roseburia hominis* A2-183, which are the only species having physically linked mec-box-like structure in non-*Staphylococcus* species (Figure 3). Our tree dissimilarity measure between SCCmec and core genes clearly indicate that SCCmec genes were undergone frequent exchange within the species. Notably, a couple of SCCmec core genes such as *orfX* and *mecR* might have been less influenced by LGT because their dissimilarity indices were similar to that of core genes (Figure 4). Due to lack of availability of diverse set of complete genomes within each *Staphylococcus* species and frequent exchange of genes within the species, it is hard to polarize LGTs happened between closely related species. Phylogenomic studies with additional diverse set of genomes of *Staphylococcus* species will provide further insights on the resistance evolution of *S. aureus*.

## Author’s contributions

S.N.-S. and J.J. designed and conceived the study. J.J. and S.M.G coordinated the study and carried out the data analysis part. S.N.-S. and J.J. drafted the manuscript. All authors gave final approval for publication.

## Acknowledgements

We thank Prof. M. Radhakrishna Pillai, the Director, and Rajiv Gandhi Centre for Biotechnology (RGCB), for the strong support to setting up the computational facility.

